# Resistance profiles and genes of *Enterobacteriaceae* from cetaceans stranded in Philippine waters from 2018-2019 provide clues on the extent of antimicrobial resistance in the marine environment

**DOI:** 10.1101/2024.12.14.628494

**Authors:** Ren Mark D. Villanueva, Jamaica Ann A. Caras, Windell L. Rivera, Maria Auxilia T. Siringan, Lemnuel V. Aragones, Marie Christine M. Obusan

## Abstract

With the premise that cetaceans are sentinels for understanding the extent of antimicrobial resistance in the marine environment, we determined the phenotypic and genotypic antibiotic resistance profiles of the *Enterobacteriaceae* isolated from cetaceans (representing twelve cetacean species) that stranded in Philippine waters from 2018-2019. The phenotypic identifications and antibiotic susceptibility profiles of the isolates were determined through VITEK 2 system while their genotypic identifications were confirmed through 16S rRNA gene sequencing. Targeted antibiotics for profiling phenotypic resistance include penicillins, cephalosporins, carbapenems, quinolones, polymyxins and folate pathway inhibitors while detected antibiotic resistance genes (ARGs) for evaluating genotypic resistance include: (1) ampicillins (*bla*_AmpC_); (2) cephalosporins (*bla*_AmpC_ *bla*_TEM_, *bla*_SHV_, and *bla*_CTX-M_); (4) carbapenem (*bla*_KPC_); (4) polymyxins (*mcr-*1) and (5) sulphonamides (*sul*1, and *sul*2). Percent resistances (% R), percent susceptibilities (% S) and multiple antibiotic resistance (MAR) index values were computed. Eighty-six *Enterobacteriaceae* were isolated from the exhaled breath condensate and swab samples of 19 stranded cetaceans. These isolates were confirmed to belong to the following genera: *Escherichia* (39.53%), *Enterobacter* (26.74%), *Klebsiella* (24.41%), *Citrobacter* (5.81%), *Morganella* (1.16%)*, Pantoea* (1.16%) and *Providencia* (1.16%). Overall, 35/86 (40.70%) of the isolates exhibited acquired resistances against cephalosporins (i.e., cefuroxime, 26/86 or 30.23%), polymyxins (i.e., colistin, 6/86 or 6.97%), folate-pathway inhibitors (i.e., trimethoprim-sulfamethoxazole,5/86 or 5.82%), ampicillin (3/86 or 3.49%), and cefoxitin (2/86 or 2.32%), while the lowest resistance (1.16% of isolates) were resistant against amoxicillin-clavulanic acid, piperacillin and imipenem. Moreover, 40.70% of the isolates were characterized as multidrug-resistant (2.33%) and extensively drug-resistant (38.37%) while 5/86 (5.81%) of the isolates had MAR indices greater than 0.2. Six out of seven (85.71%) of the targeted ARGs responsible for the resistance types for ampicillins, cephalosporins, polymyxins and sulphonamides (i.e., *bla*_AmpC_, *bla*_SHV_ *bla*_TEM,_ *mcr-1, sul*1 and *sul*2, respectively) were detected in 48.57% of isolates. Antibiotic susceptibility testing revealed that a considerable portion of the isolates exhibited acquired resistance to selected antibiotics and were categorized as multidrug-resistant (MDR) or extremely drug-resistant (XDR). As for genotypic resistance, six out of seven target antibiotic resistance genes (ARGs) responsible for resistance to ampicillins, cephalosporins, polymyxins, and sulfonamides were detected in nearly half of the isolates with acquired resistance. Considering the habitat ranges of the source animals, this indicates the extent of reach of antibiotics and/or ARGs in the marine environment, and pelagic migratory cetaceans may play an important role in their dissemination.

## Introduction

The World Health Organization (WHO) calls for increased efforts to address the threat of bacterial resistance to antibiotics that are both used in human and veterinary medicine [1]. Antimicrobial resistance (AMR) is a rising global healthcare concern for human health, causing an estimate of 700,000 annual mortality which, if not addressed, is projected to increase to 10 million mortality per year by 2050 [2]. It is also a critical concern in agriculture for the maintenance of livestock animal health [3]. Given the foregoing facts, it is essential to improve our understanding of AMR in humans and animals. In general, the attention to AMR in animals is biased towards domestic species, with less efforts for its investigation in wildlife. Nevertheless, there is evidence of the prevalence of AMR in wildlife species and natural environments [3–5].

The available literature on bacteria that were isolated from marine mammals worldwide support the significance of investigating *Enterobacteriaceae* and their antibiotic resistance genes (ARGs). *Enterobacteriaceae* accounts for a significant proportion of human infections worldwide, and is a major concern [6]. Members of this bacterial group has reported resistances against penicillins, cephalosporins, cephamycins, and carbapenems conferred by β-lactamases [7]. In 2017, WHO listed *Enterobacteriaceae* as a critical priority for antibiotic resistance surveillance [1; 8]. Studies reporting AMR in marine mammals are fewer compared to terrestrial animals, and more studies are needed to understand the dissemination of resistant bacteria in the marine environment [9]. In the Philippines, Obusan et al. (2018) provided the initial report on antibiotic susceptibility of bacteria isolated from Philippine cetaceans, wherein more than half of these bacteria (n=14) had single or multiple resistances to a selection of antibiotics [10]. Elsewhere, Delport et al. (2015), initially reported the ARGs (i.e. streptomycin-spectinomycin (*aadA1*) and trimethoprim (*dfrA17, and dfrB4*) present among marine mammals particularly in endangered Australian sea lions (*Neophoca cinerea*) [11]. More recently, Vale et al. (2021) reported the detection of *β* -lactamase encoding genes in *Escherichia coli* from rescued seals, *bla*_OXA−1_ and *bla*_TEM−1_) while Grünzweil et al., (2021) reported *bla*_CMY_ as the predominant *β*-lactamase gene among bacterial isolates that exhibited MDR phenotype, followed by other *β-*lactamase genes (*bla*_TEM-1,_ *bla*_SHV-33,_ *bla*_CTX-M-15_, *bla*_OXA-15_, *bla*_SHV-11_ and *bla*_DHA-11_) [12–13]. On the other hand, the most prevalent non-*β*-lactamase gene was *sul*2, followed by *str*A, *stra*B and *tet*(A) [13].

The antibiotic resistance patterns of bacteria isolated from cetaceans can provide clues on common antibiotic classes and ARGs circulating in their habitats in relation to anthropogenic activities such as the unregulated use of antibiotics in aquaculture [14]. One study in the Philippines reported the presence of sulfamethoxazole (SMX) resistance genes (i.e., *sulI* and *sulII* in culturable bacteria, *sulIII* in unculturable bacteria) in bacterial communities of Laguna Lake, Pasig River, and Manila Bay [15].

This study aimed to (1) isolate *Enterobacteriaceae* from cetaceans that stranded in the country from 2018-2019; (2) determine the antibiotic susceptibility profiles of the isolates; (3) detect ARGs of the isolates; and (4) analyze cetacean strandings in association with the occurrence and patterns of antibiotic resistance as well as ARGs in the isolates.

## Materials and Methods

### Responding to cetacean stranding events

Cetacean stranding events that occurred in the Philippines from January 2018-June 2019 were responded in collaboration with Philippine Marine Mammal Stranding Network (PMMSN) and Bureau of Fisheries and Aquatic Resources (BFAR) of the Department of Agriculture (DA). PMMSN is an active partner of the Microbial Ecology of Terrestrial and Aquatic Systems Laboratory (METAS-Aquatic), University of the Philippines-Institute of Biology (UP-IB), and Marine Mammal Research and Stranding Laboratory (MMRSL), University of the Philippines-Institute of Environmental Science and Meteorology (UP-IESM); the network cooperates in a scheme for specimen collection and transport to laboratories.

Stranded cetaceans were characterized in terms of: (1) species; (2) sex (based on genital and/or mammary slits); (3) length (tip of the snout to the tip of the tail or notch of the flukes); (4) age class (inferred from length based on species); (5) body condition (based on the criteria provided by Read & Murray [16]; (6) stranding type (e.g., single or mass; live or dead); (7) stranding site; and (8) stranding season (based on Wang, 2006) [17]. Biological material collection was performed considering the (1) disposition of the animal (e.g., alive, or dead); (2) physical preservation of the animal (i.e., based on the expanded version of the Code system established by the Smithsonian Institution’s Marine Mammal Events Program [18]; and (3) conditions in the stranding site (e.g., handling limitations that may result to sample contamination). The following physical preservation codes were used: Code 1-live animal; Code 2 – fresh (carcass in good condition); Code 3-fair (decomposed, but organs basically intact); Code 4-poor (advanced decomposition); and Code 5 – mummified or skeletal remains.

### Isolation of bacteria

Bacterial isolates were derived from the biological samples collected by the members (e.g. veterinarians) of PMMSN and DA-BFAR who were trained in medical aspects of marine mammal response. Samples from routine and non-routine sites using sterile cotton or synthetic swabs in a transport medium (e.g., Amie’s (HiMedia Laboratories, India) [19]. For routine sites, swab samples were collected from the blowhole and rectum of live and freshly dead cetaceans [20–23]. In the case of live stranders, the swab was inserted into the blowhole during a breath, and the swab was swiped down to about 1-3 inches. Whenever applicable, liquid droplets and exhaled breath were also collected by lowering a sterile dish. Non-routine sites for swabbing included lesions, organs, and thoracic fluid. Swab samples were stored in coolers with ice gel packs (4°C −7°C) and shipped to the laboratory within 24 h for immediate processing [24].

From each transport medium, 10 mL were mixed with 90 mL Tryptone Soy Broth (TSB) (HiMedia Laboratories, India) and incubated for 24 h at 37°C. If growth was observed, indicated by turbidity, 1 mL of the TSB culture was transferred into 99 mL MacConkey Broth (HiMedia Laboratories, India) and incubated for 48 h at 37°C to enhance the growth of members of *Enterobacteriaceae*. After enrichment, isolation of enteric bacteria was done by streaking on MacConkey agar (HiMedia Laboratories, India) plates and incubated at 37°C for 72 h. Distinct colonies on MacConkey agar plates were streaked into MacConkey agar slants [25]. A total of 86 Gram-negative enteric bacteria were isolated from stranded cetaceans.

### Phenotypic and genotypic identification of isolates

Pure cultures of 86 putative enteric bacterial isolates (n=86) were Gram-stained following Hucker’s method [26] and were examined under the microscope (Olympus CX23, Japan). Isolates exhibited Gram-negative rod-shaped cells were further examined. Using the VITEK 2 system and VITEK GN cards, (bioMerieux, Inc. France), putative enteric bacterial isolates were confirmed as members of *Enterobacteriaceae* based on the pertinent biochemical reactions which include carbon utilization and enzymatic activities [27].

For molecular identification, extraction of bacterial DNA was done using a commercially available kit (GF-1 Bacterial DNA Extraction Kit) following manufacturer’s instructions. Primers 27F (5’-AGAGTTTGATCCTGGCTCAG-3’) and 1541R (5’-AAGGAGGTGATCCANCCRCA-3’) were used for the amplification of universal 16S rRNA bacterial gene [28]. Reactions were performed in 25 μl volume with the following concentrations of components: 1X PCR Master Mix (Promega: contains *Taq* DNA polymerase, dNTPs, MgCl_2_), 1.0 µM assigned primers, 1.5-3.0 µL of 40-50 ng of DNA template and nuclease free water adjusted accordingly [28]. The optimized PCR conditions used in the identification include the following parameters: initial denaturation at 95°C for 2 min, followed by 30 cycles of 94°C for 30 s, 55°C for 30 s, and 72°C for 30 s and final extension from 7 min at 72°C. Negative controls (DNA-free templates) were included. Electrophoresis of PCR products in TAE (Tris-acetate-EDTA) buffer were performed on 1% agarose gels at 100 V with DNA ladder. Following the electrophoresis, the gels were stained using a nucleic acid dye and viewed under UV light to detect target DNA band. PCR-positive samples were then sent to Macrogen, South Korea for purification and sequencing. Molecular analyses were performed using software programs (e.g., STADEN package; BioEdit Sequence Alignment Editor version 7.0.5.3) [29–30] for assembling, editing and aligning of sequence. The sequence alignment was analyzed by NCBI Basic Local Alignment Nucleotide (BLAST) tool available at https://blast.ncbi.nlm.nih.gov/Blast.cgi?PROGRAM=blastn&PAGE_TYPE=BlastSearch&LINK_LOC=blasthome, using the parameters “highly similar sequences (megablast)”.

### Determination of antibiotic susceptibility profiles

The susceptibility profiles of the isolates from cetaceans that stranded from 2018-2019 were determined through VITEK 2 automated system (bioMerieux, France), at the Pathogen-Host-Environment Interactions Research Laboratory (PHEIRL), UP-IB, in accordance with the manufacturer’s instructions. The system uses AST-N261 cards that contain the following antibiotics as dehydrated substances in various concentrations; amikacin, amoxicillin-clavulanic acid, ampicillin, cefepime, cefoxitin, ceftazidime, ceftriaxone, cefuroxime, ciprofloxacin, colistin, ertapenem, extended spectrum-β-lactamase, gentamicin, imipinem, meropenem, piperacillin-tazobactam and trimethoprim-sulfamethoxazole. The susceptibility profiles of the bacteria were used as reference for the detection of target ARGs.

### Detection of ARGs

Bacterial isolates that exhibited antibiotic resistance phenotypes were subjected to the detection of target ARGs (Table 1). Nine representative AMR genes responsible for the resistance phenotype of five antimicrobial categories were detected: *bla*_AmpC_, encoding for a broader spectrum spectrum of resistance, especially when present along with other classes of ESBLs; *bla*_AmpC_ *bla*_CTX-M,_ *bla*_TEM,_ and *bla*_SHV_, encoding for the cephalosporin resistance; *qnr*B, and *anr*S, encoding for the quinolone resistance; *mcr-1*, for the polymyxin resistance; and *sul*1 and *sul*2 for the sulfonamide resistance [31–35]. These targets include those proven to confer resistance mechanisms for antibiotics used in the susceptibility assay. Prior studies reported the detection of these ARGs in heavily impacted water bodies such as aquaculture systems and sewage discharge river, as well as in relatively pristine aquatic environments [36].

**Table 1.**
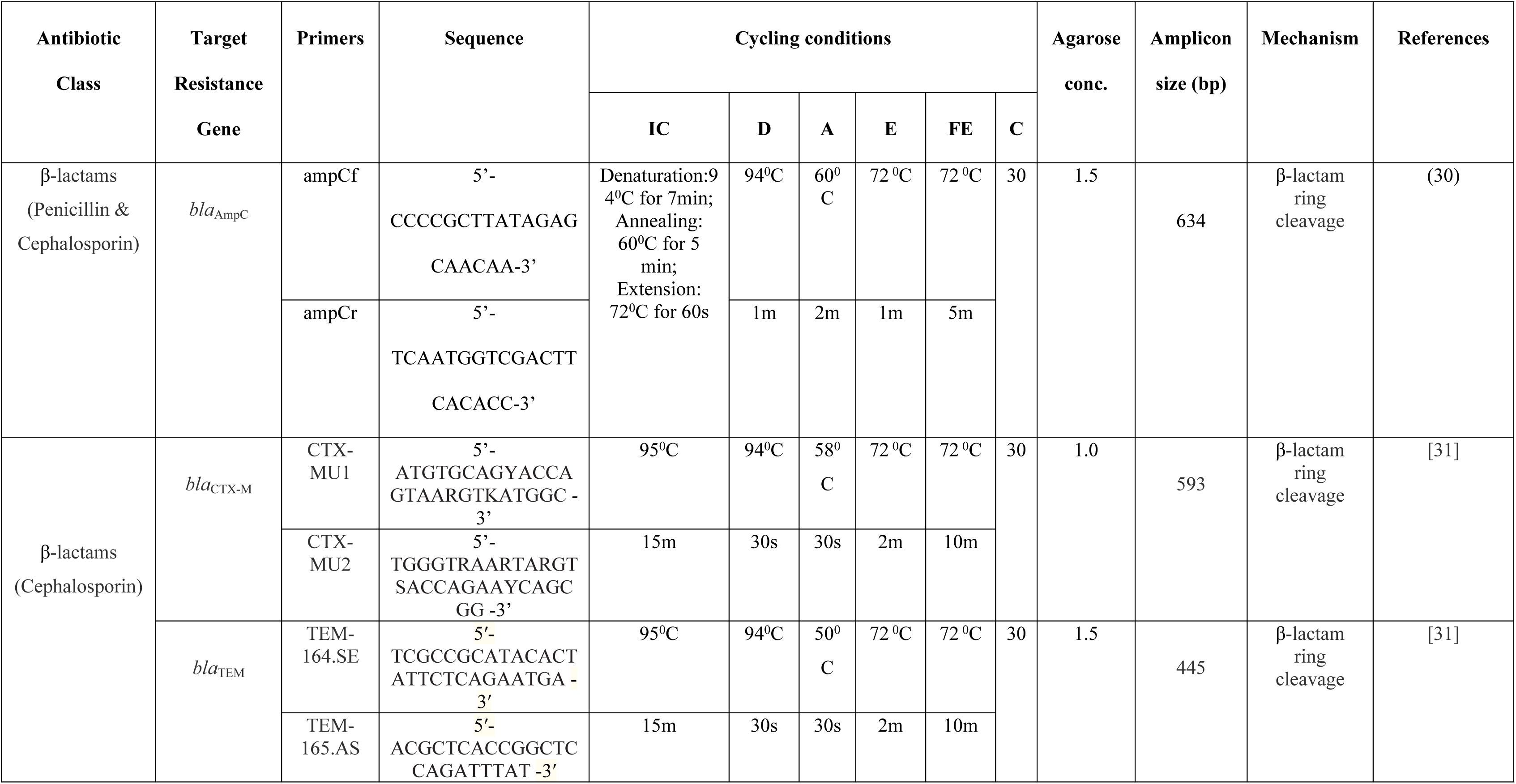
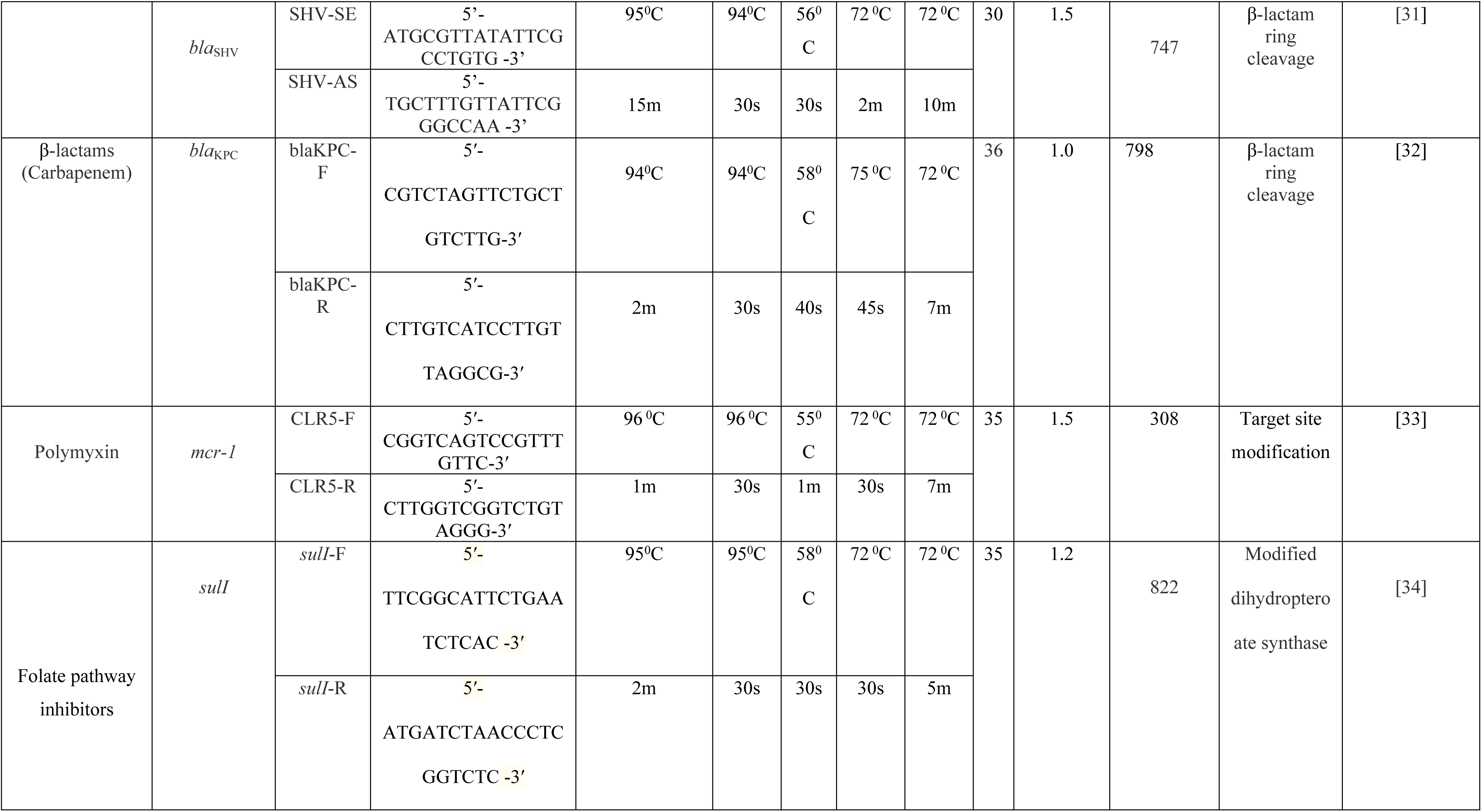
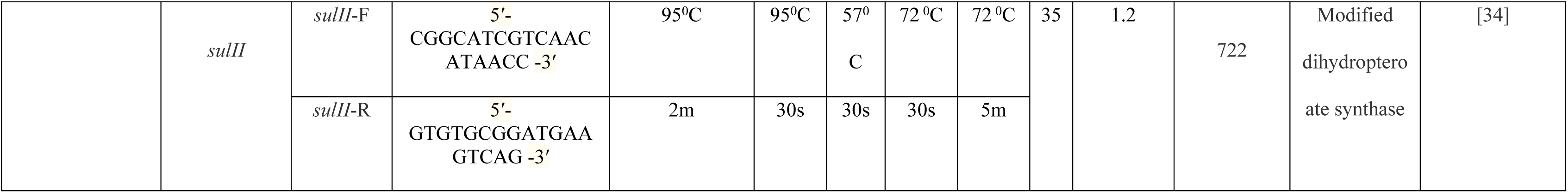
List of target ARGs.

PCR reactions were carried out in a 25µl reaction mixture with the following concentration of components: 1X PCR Master Mix (Promega: contains *Taq* DNA polymerase, dNTPs, MgCl_2_), 1.0 µM assigned primers, 1.5-3.0 µL of 40-50 ng of DNA template and nuclease free water adjusted accordingly [28] and conditions specific to each target gene. Positive controls included either DNA from antibiotic resistant isolates (ATCC controls *Klebsiella pneumoniae* BAA-1705, *Escherichia coli* 35218, selected antibiotic-resistant isolates from microbial culture collection of IB-MCCAquaS, UP-nd negative controls exclude the DNA template. Electrophoresis of PCR products in TAE buffer were performed on 1-1.5% agarose gels at 100 V with DNA ladder. Following the electrophoresis, the gels were stained using a nucleic acid dye and viewed through UV light exposure to detect target DNA band.

### Data analysis

Percent resistances (% R) and percent susceptibilities (% S) were computed per bacterial isolate and antibiotic, and cetacean individual. Multiple antibiotic resistance (MAR) index values were computed using the formula: number of resistant antibiotics / total number of antibiotics tested [37] MAR indices greater than 0.2 were interpreted to come from sources where antibiotics were often used [8; 37]. In addition, MAR isolates were interpreted as those that have acquired resistance to three or more antibiotic classes [38]. Each isolate was characterized based on the presence/absence of each resistance phenotype and target resistance gene. Multiple AMR is described as phenotypic resistance to more than two antimicrobials while multiple resistance gene pattern is described as presence of more than two ARGs in an isolate. Isolates with intermediate MIC were considered as susceptible [39].

## Results

### Stranded cetaceans

*Enterobacteriaceae* were isolated from 19 stranded cetaceans representing 12 species (*Balaenoptera edeni, B. omurai, Feresa attenuata, Grampus griseus, Kogia breviceps, Lagenodeplhis hosei, Mesoplodon densirostris, Peponocephala electra, Stennela attenuata, S. coeruleoalba, Tursiops aduncus,* and *T. truncatus*). The stranding events of these cetaceans were responded in different regions of the Philippines, mostly in Regions V, IX and X, and the rest in Region I, Region II, Region IV-B, Region XI and Region XII (Fig 1). Majority of the cetaceans (10) were female, while there are 7 males; the sex of the remaining two cetaceans was unknown (S1 Table).

**Fig. 1.**
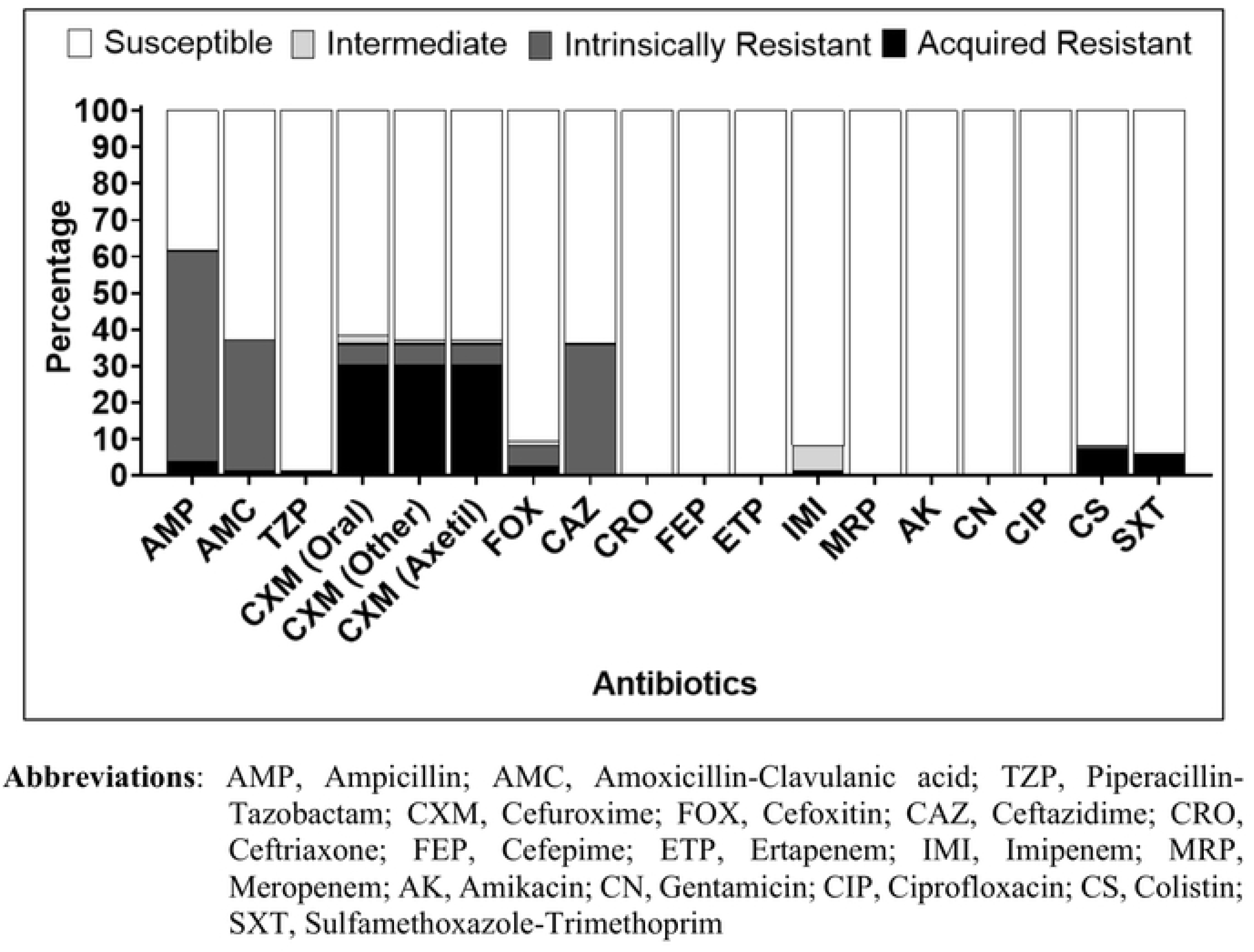
Cetacean stranding sites (Nineteen (19) individuals belonging to twelve (12) cetacean species that stranded in Philippine waters from February 2018 to May 2019. (Cetacean stranders were sampled for bacterial isolation) (Reprinted Wikimedia Commons, the free media repository, 2009).

### *Enterobacteriaceae* isolated from stranded cetaceans

Among the *Enterobacteriaceae* isolates (n-86), *Escherichia* sp. is the predominant genus (n=34 or 39.53%), followed by *Enterobacter* sp. (n=23 or 26.74%), *Klebsiella* sp. (n=21 or 24.41%), and *Citrobacter* sp. (n=5 or 5.81%). The remaining three isolates were identified as *Morganella morganii, Pantoea agglomerans* and *Providencia alcalifaciens* (Fig 2 and S1 Table).

**Fig. 2.**
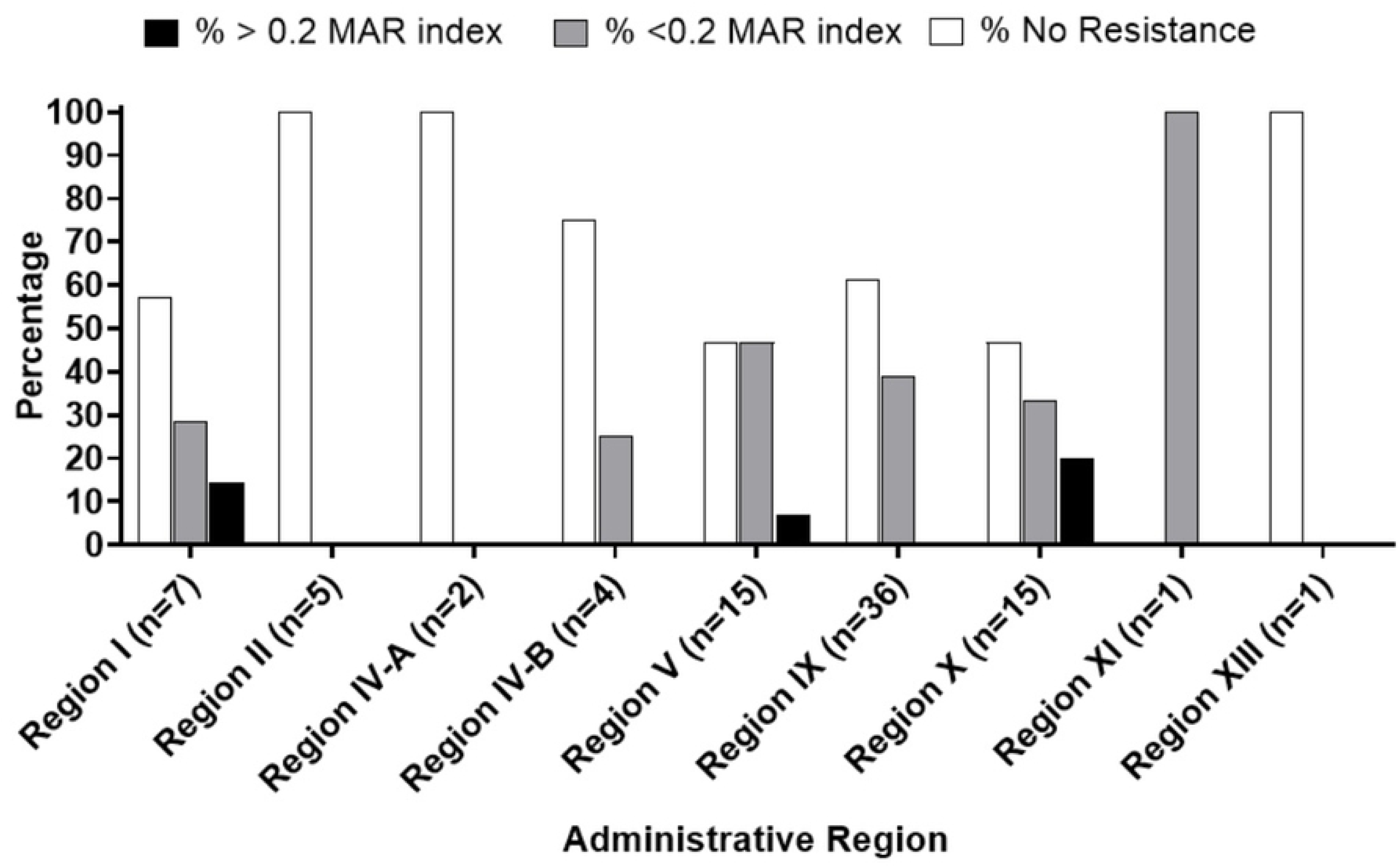
Distribution of *Enterobacteriaceae* (n=86) isolated from the stranded cetaceans. Phenotypic antibiotic resistance patterns of isolated *Enterobacteriaceae*.

The resistance of *Enterobacteriaceae* isolates against a spectrum of 18 antibiotics from different classes was analyzed and these microorganisms exhibited variable antibiotic susceptibility patterns (S2 Table). Based on the susceptibility patterns, aminoglycosides (amikacin and gentamicin), carbapenems (ertapenem and meropenem), cephalosporins (ceftazidime, ceftriaxone and cefepime) and quinolones (ciprofloxacin) were the most effective antibiotics. On the other hand, all isolates exhibited the highest acquired resistance phenotype against cephalosporins: 26/86 (30.23%) for cefuroxime (oral, axetil and other). The following are the percentages of acquired resistance phenotype against the remaining antibiotics: colistin 6/86 (6.97%), followed by trimethoprim-sulfamethoxazole 5/86 (5.82%), ampicillin 3/86 (3.49%) and cefoxitin 2.86 (2.32%), while the lowest resistance (1.16% of isolates) was observed against amoxicillin-clavulanic acid, piperacillin, and imipenem. Lastly, none of the isolates were positive for extended spectrum *β*-lactamases (ESBL) resistance, i.e., isolate that confers resistance to most *β*-lactam antibiotics used in this study (Fig 3 & S2 Table).

**Fig. 3.**
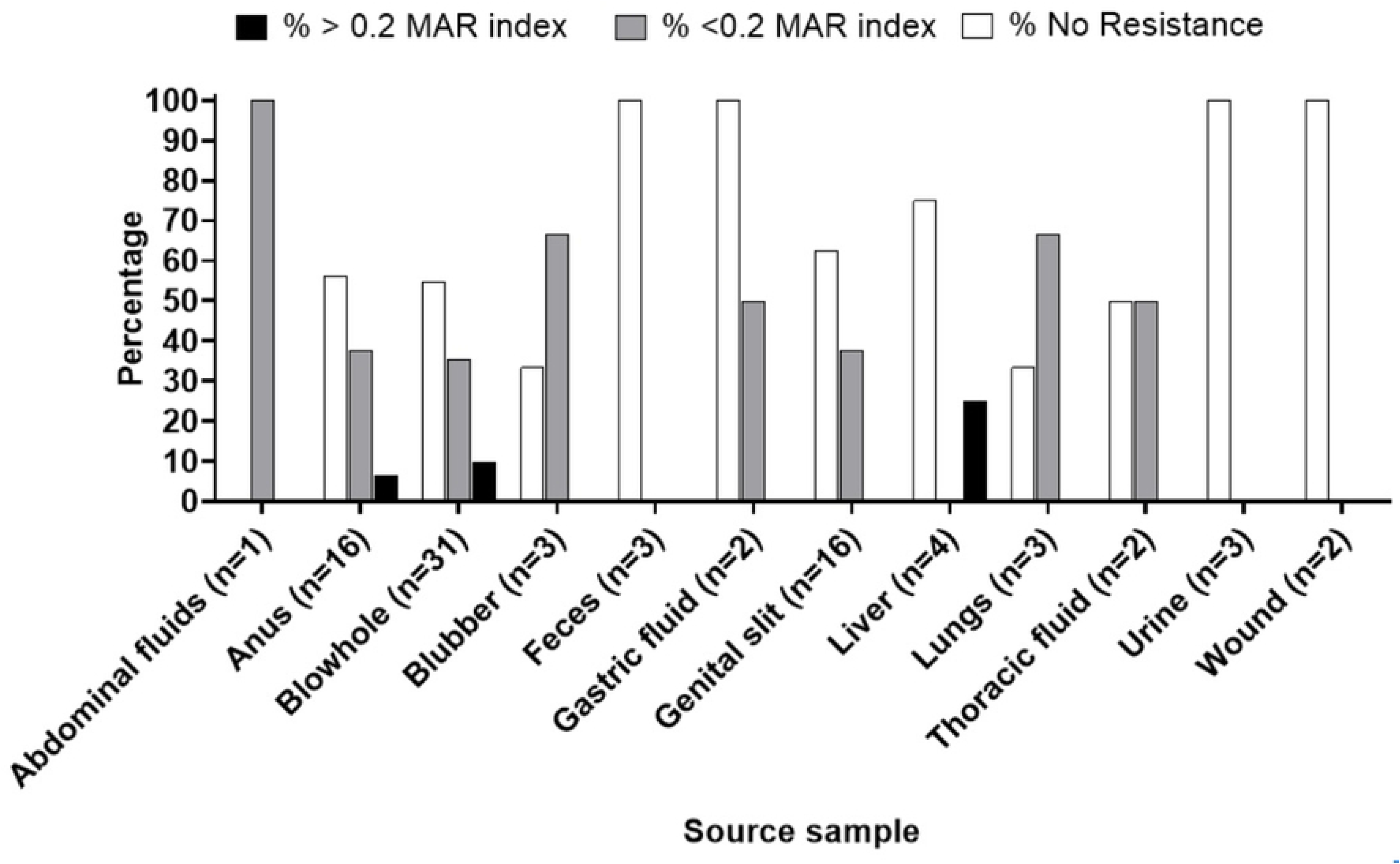
% Resistances of *Enterobacteriaceae* isolated from stranded cetaceans.

About 40.7% of the isolates (35/86) exhibited acquired resistance phenotype against 18 tested antibiotics were determined. The highest MAR index value (0.27) was observed in isolates 18-WW, 18-M3, 18-K4 & 18-Q4, which were molecularly identified as *Enterobacter cloacae* complex, were all resistant to 4 of the 15 tested antibiotics. Followed by isolate 18-XX (named as *Escherichia coli*) which has the next highest MAR index value, that is, 0.22, was shown to be resistant to 4 antibiotics (Fig 4 & S1 Table). In line with this, the MAR index values of these isolates are higher than 0.2, which is suggested to be a characteristic of point-source pollution (Fig 4 & S3 Table) [9; 32].

**Fig. 4.**
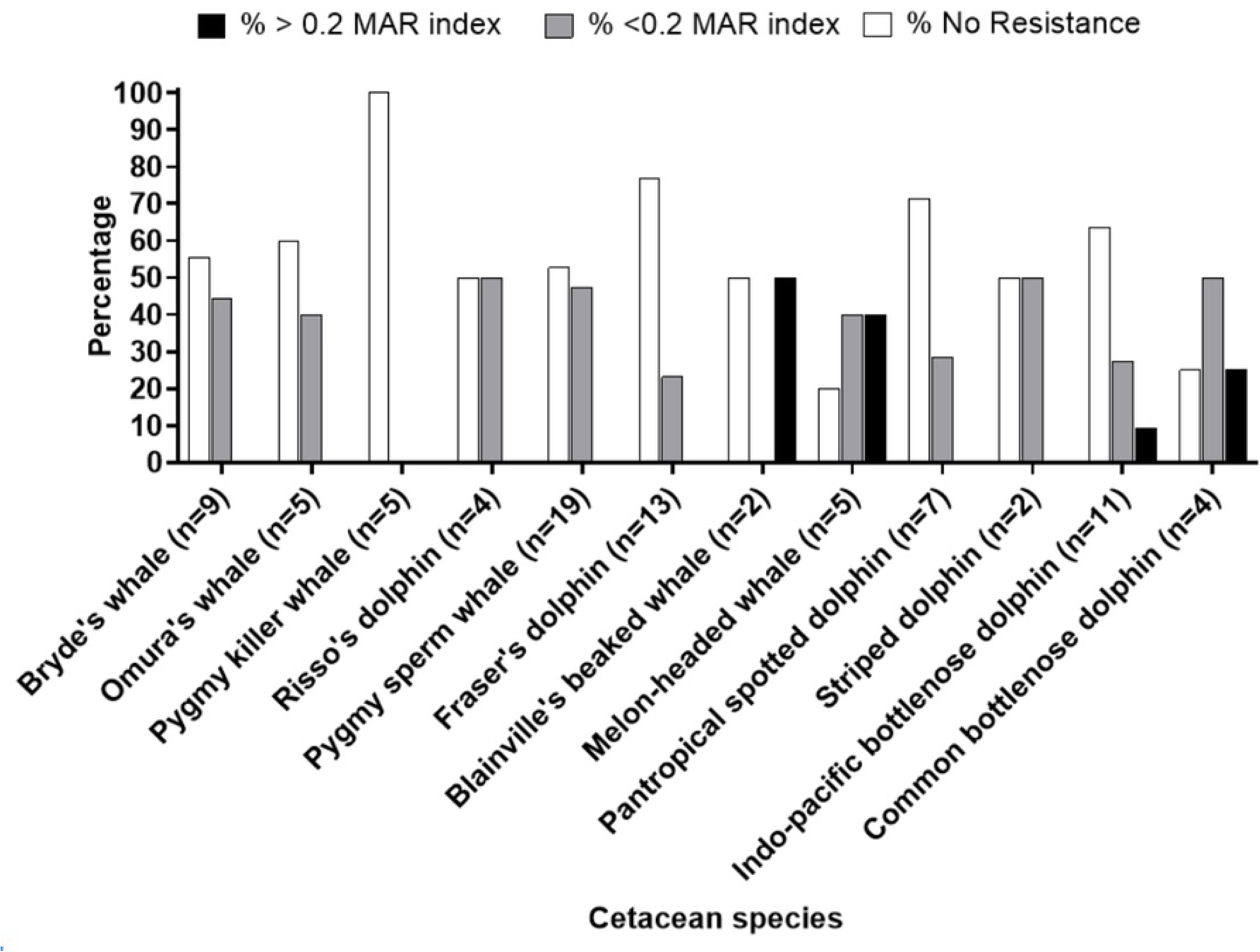
Multiple antibiotic resistance (MAR) indices of selected *Enterobacteriaceae* isolated from the stranded cetaceans.

Isolates belonging to *Enterobacteriaceae* with MAR indices of > 0.2 were derived from cetacean species that are known to inhabit coastal zone (e.g., *Tursiops truncatus* (common bottlenose dolphins)) and pelagic zone (e.g. *Mesoplodon densirostris* (Blainville’s beaked whale, *Tursiops aduncus* (Indo-pacific bottlenose dolphin) & *Peponocephala electra* (Melon-headed whale)). Moreover, bacteria with a MAR index of > 0.2 were isolated from internal body sites (1/86) (i.e., isolate 18-Q4 (*Enterobacter cloacae*) (Fig 5 & S4 Table).

**Fig. 5.**
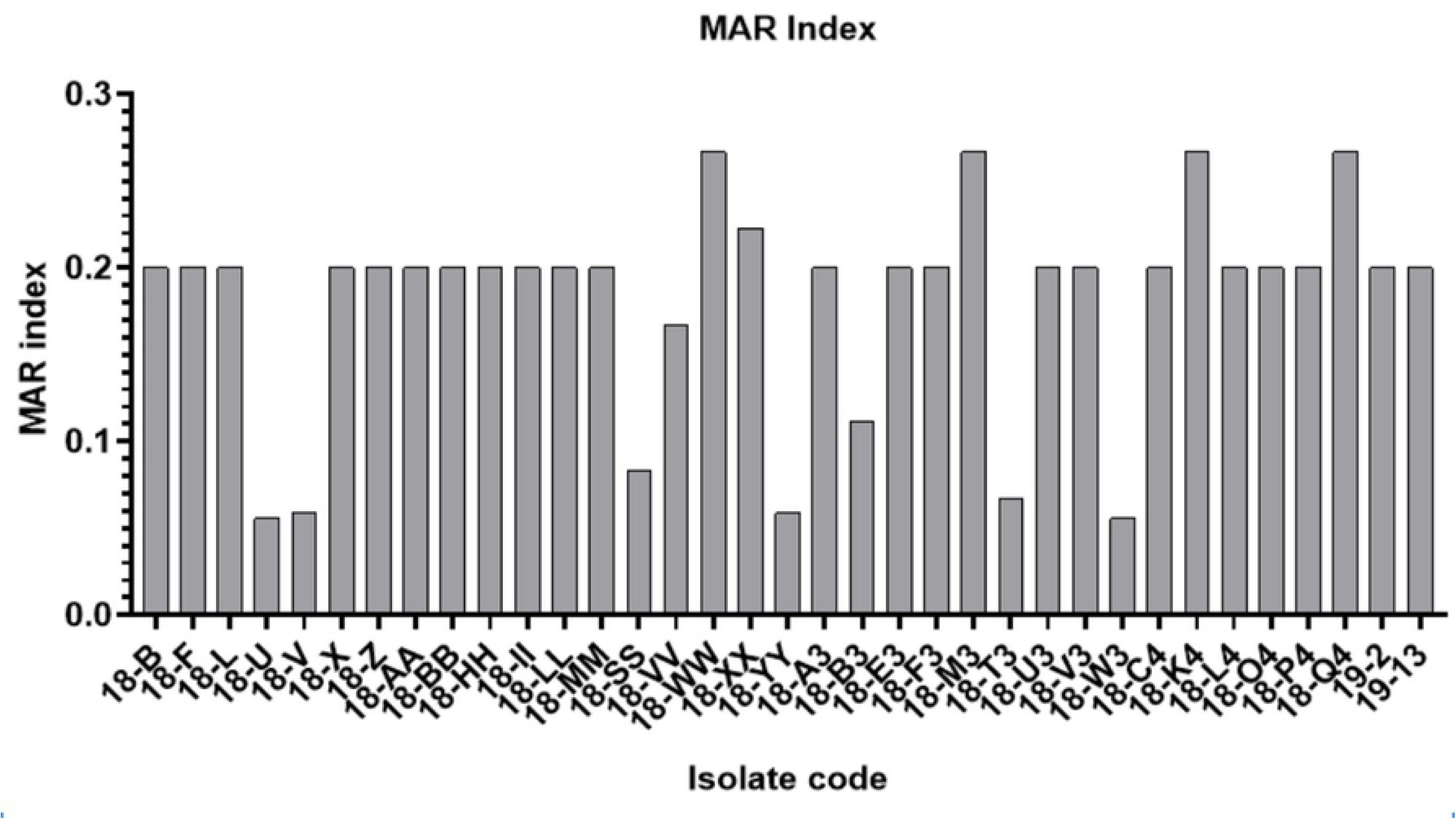
Prevalence of multiple antibiotic resistances among *Enterobacteriaceae* isolated from different cetacean species.

Bacteria (4/86) isolated from body sites (i.e., blowhole and anus) that are in contact with the environment have > 0.2 MAR indices as compared to internal body sites such as those isolated from the anus (i.e., of isolates 18-K4 (*Enterobacter cloacae*) and blowhole (i.e., isolates 18-WW & 18-M3 (*Enterobacter cloacae*) and 18-XX (*Escherichia coli*) (Fig 6 & S5 Table).

**Fig. 6.**
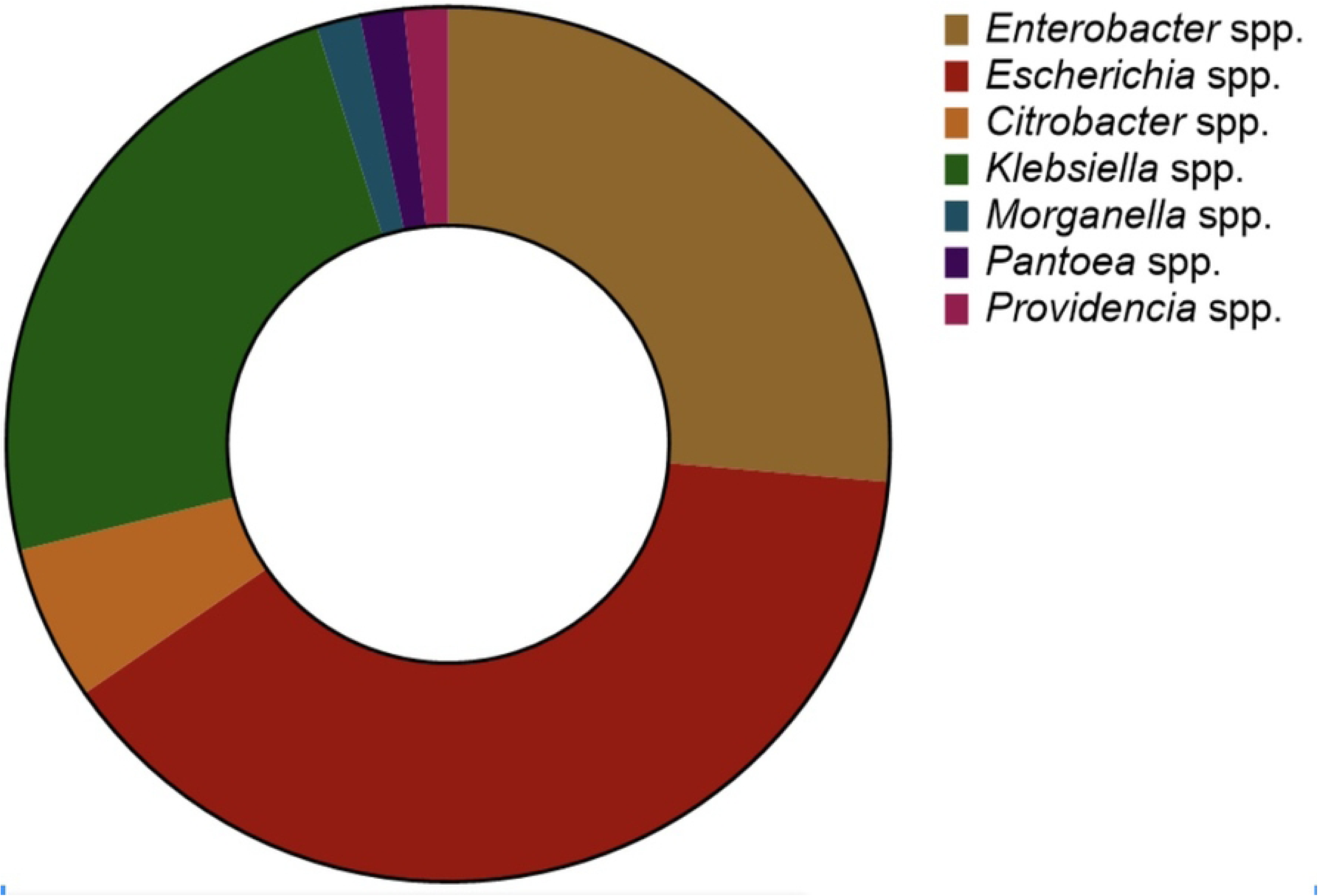
Prevalence of multiple antibiotic resistances among *Enterobacteriaceae* isolated from different source samples.

In terms of stranding sites, percentages of bacteria isolated from cetaceans with MAR index values that exceed 0.2 were mostly coming from Region X (20.00%), Region I (14.29%) and Region V (6.67%) (Fig 7 & S6 Table).

**Fig. 7.**
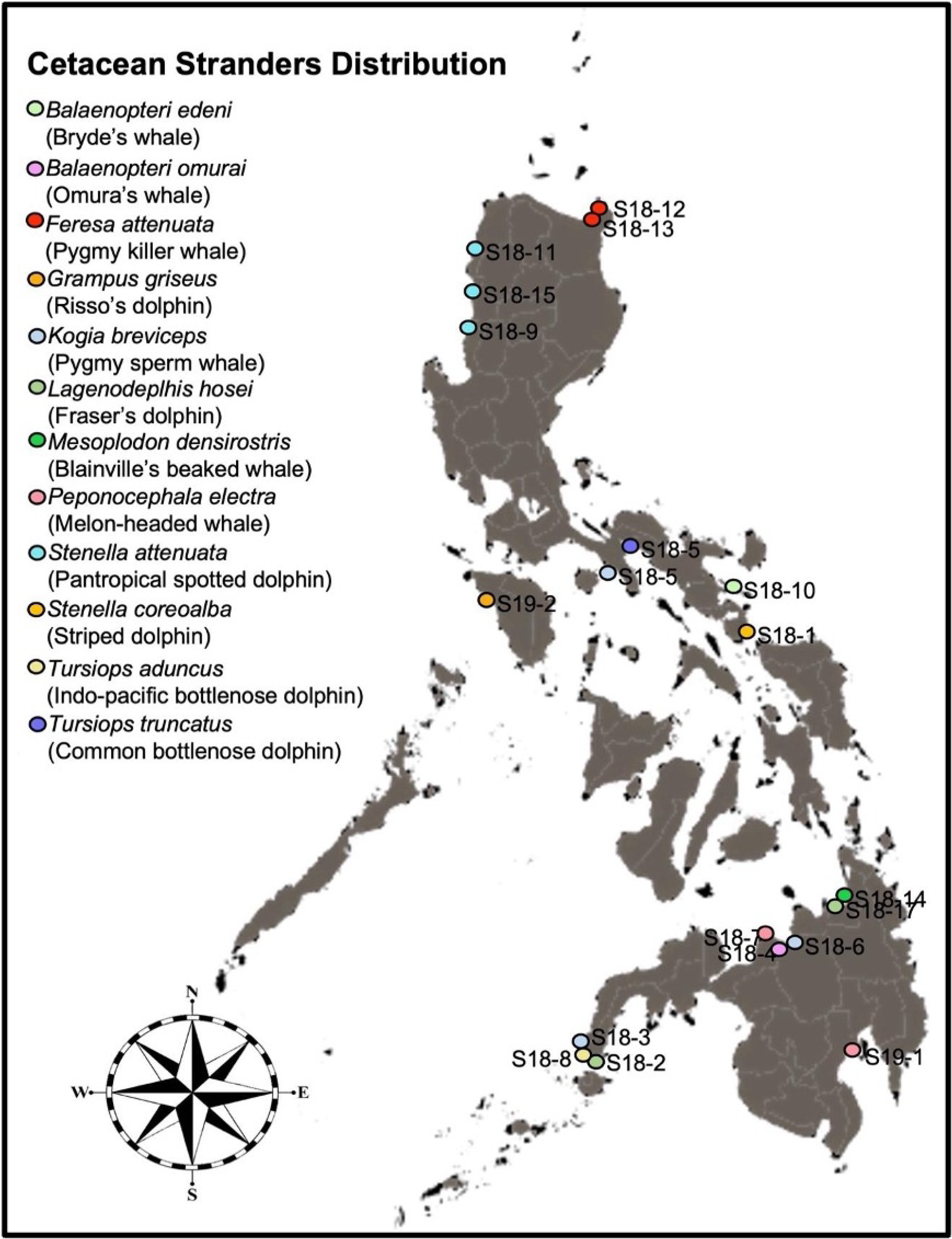
Prevalence of multiple antibiotic resistances among *Enterobacteriaceae* isolated from different administrative regions.

### Multidrug-resistant (MDR) *Enterobacteriaceae*

Multidrug-resistant (MDR) isolates were further categorized following the criteria of Magiorakos et al. (2012): a) MDR are bacteria that have acquired resistance to at least one antimicrobial agent in three or more antimicrobial classes; b) extensively drug-resistant (XDR) are bacteria that have acquired resistance to at least one to two antimicrobial classes; and c) pandrug-resistant (PDR) are bacteria that have acquired resistance in all antimicrobial classes [38]. Two isolates (i.e., isolates 18-VV and 18-XX identified as *E. coli*) were categorized as MDR, while 33 of which showed acquired resistance phenotype were categorized XDR, suggesting the high incidence of multidrug resistance among the cetacean-derived enteric bacterial isolates (40.70%). There were no isolates categorized as PDR (Table 2 & S3 Table).

**Table 2.** Classification and percentage of acquired antibiotic resistances of *Enterobacteriaceae* isolated from cetaceans.

| Organism | Total isolates | MDR | XDR | PDR |
| --- | --- | --- | --- | --- |
| *Citrobacter* sp. | 5 | 0 (0.00%) | 1 (20.00%) | 0 (0.00%) |
| *Enterobacter* sp. | 25 | 0 (0.00%) | 25 (100.00%) | 0 (0.00%) |
| *Escherichia* sp. | 34 | 2 (5.88%) | 2 (5.88%) | 0 (0.00%) |
| *Klebsiella* sp. | 19 | 0 (0.00%) | 3 (15.79%) | 0 (0.00%) |
| *Morganella* sp | 1 | 0 (0.00%) | 0 (0.00%) | 0 (0.00%) |
| *Pantoea* sp. | 1 | 0 (0.00%) | 1 (100.00%) | 0 (0.00%) |
| *Providencia* sp. | 1 | 0 (0.00%) | 1 (100.00%) | 0 (0.00%) |
| **Total** | **31** | **2 (2.33%)** | **33 (38.37%)** | **0 (0.00%)** |

### Prevalence of ARGs from MDR and XDR isolates

ARGs were detected from the isolates with acquired resistance phenotypes. All four isolates (100.00%) with acquired penicillin resistance phenotype possess *bla*_AmpC_; isolates 18-VV (*E. coli*), 18-SS (*C. freundii*), 18-XX (*E. coli*) and 18-B3 (*E. coli*). ARGs responsible for cephalosporin resistance phenotype were also detected; *bla*_SHV_ in isolates 18-F (*E. cloacae*), 18-L (*E. cloacae*), 18-V (*P. agglomerans*), 18-Z (*E. cloacae* complex), 18-AA (*E. cloacae*), 18-LL (*E. cloacae*) and 18-MM (*K. aerogenes*). Aside from *bla*_SHV_, *bla*_TEM_ was also detected in isolates 18-L (*E. cloacae*), 18-V (*P. agglomerans*), 18-WW (*E. cloacae* subsp. dissolvens), 18-XX (*E. coli*) and 18-A3 (*E. cloacae*). Other cephalosporin resistance gene, specifically *bla*_CTX-M_ was not detected. One carbapenem-resistant isolate, 18-T3 (*E cloacae* complex), does not possess any carbapenem-resistance gene (i.e., *bla*_KPC_). Moreover, four out of five sulfonamide-resistant isolates (i.e., 18-VV (*E. coli*), 18-YY (*K. pneumoniae*), 18-B3 (*E. coli*) and 18-W3 (*E. coli*) possess *sul*2 gene, and three of these isolates (i.e., 18-VV (*E. coli*), 18-YY (*K. pneumoniae*) and 18-B3 (*E. coli*)) also possessed *sul*1 gene. One of the genes responsible for the polymyxin resistance phenotype, *mcr-1*, was detected in 18-K4 (*E. cloacae* subsp. dissolvens) and 18-Q4 (*E. cloacae*). Overall, multiple resistance genes (i.e., presence of more than two resistance genes) were 18-VV and 18-B3 that were both molecular identified as *E. coli* both possess *bla*_AmpC,_ *sul*1 and *sul*2 (Table 3 & S1-S8 Figs).

**Table 3.** Prevalence of antibiotic resistance genes (ARGs) from *Enterobacteriaceae* with acquired antibiotic resistances.

| Isolate Code | Molecular Identity | Penicillin Resistance gene | Cephalosporin resistance genes |  |  | Carbapenem Resistance gene | Polymyxin Resistance gene | Sulfonamide Resistance genes |  |
| --- | --- | --- | --- | --- | --- | --- | --- | --- | --- |
|  |  | *bla*<sub>AmpC</sub> | *bla*<sub>CTX-M</sub> | *bla*<sub>SHV</sub> | *bla*<sub>TEM</sub> | *bla*<sub>KPC</sub> | *mcr-1* | *sul1* | *sul2* |
| 18-B | *Enterobacter cloacae* complex |  | - | - | - |  |  |  |  |
| 18-F | *Enterobacter cloacae* |  | - | + | - |  |  |  |  |
| 18-L | *Enterobacter cloacae* |  | - | + | + |  |  |  |  |
| 18-U | *Providencia alcalifaciens* |  |  |  |  |  | - |  |  |
| 18-V | *Pantoea agglomerans* |  | - | + | + |  |  |  |  |
| 18-X | *Enterobacter cloacae* |  | - | - | - |  |  |  |  |
| 18-Z | *Enterobacter cloacae* complex |  | - | + | - |  |  |  |  |
| 18-AA | *Enterobacter cloacae* |  | - | + | - |  |  |  |  |
| 18-BB | *Enterobacter cloacae* |  | - | - | - |  |  |  |  |
| 18-HH | *Enterobacter cloacae* |  | - | - | - |  |  |  |  |
| 18-II | *Enterobacter cloacae* |  | - | - | - |  |  |  |  |
| 18-LL | *Enterobacter cloacae* |  | - | + | - |  |  |  |  |
| 18-MM | *Klebsiella aerogenes* |  | - | + | - |  |  |  |  |
| 18-VV | *Escherichia coli* | + |  |  |  |  | - | + | + |
| 18-WW | *Enterobacter cloacae* subsp. dissolvens |  | - | - | + |  |  | - | - |
| 18-SS | *Citrobacter freundii* | + |  |  |  |  |  |  |  |
| 18-XX | *Escherichia coli* | + | - | - | + |  | - |  |  |
| 18-YY | *Klebsiella pneumoniae* |  |  |  |  |  |  | + | + |
| 18-A3 | *Enterobacter cloacae* |  | - | - | + |  |  |  |  |
| 18-B3 | *Escherichia coli* | + |  |  |  |  |  | + | + |
| 18-E3 | *Enterobacter aerogenes* |  | - | - | - |  |  |  |  |
| 18-F3 | *Enterobacter aerogenes* |  | - | - | - |  |  |  |  |
| 18-M3 | *Enterobacter cloacae* |  | - | - | - |  | - |  |  |
| 18-T3 | *Enterobacter cloacae*<br>complex |  |  |  |  | - |  |  |  |
| 18-U3 | *Enterobacter cloacae*<br>complex |  | - | - | - |  |  |  |  |
| 18-V3 | *Klebsiella aerogenes* |  | - | - | - |  |  |  |  |
| 18-W3 | *Escherichia coli* |  |  |  |  |  |  | - | + |
| 18-C4 | *Enterobacter cloacae* |  | - | - | - |  |  |  |  |
| 18-K4 | *Enterobacter cloacae*<br>subsp. dissolvens |  | - | - | - |  | + |  |  |
| 18-L4 | *Enterobacter cloacae*<br>subsp. dissolvens |  | - | - | - |  |  |  |  |
| 18-O4 | *Enterobacter*<br>*hormaechei* |  | - | - | - |  |  |  |  |
| 18-P4 | *Enterobacter* sp. |  | - | - | - |  |  |  |  |
| 18-Q4 | *Enterobacter cloacae* |  | - | - | - |  | + |  |  |
| 19-2 | *Enterobacter mori* |  | - | - | - |  |  |  |  |
| 19-13 | *Enterobacter* sp. |  | - | - | - |  |  |  |  |
| **Prevalence of antibiotic**<br>**resistance genes** |  | 100.00% | 0.00% | 25.00% | 17.86% | 0.00% | 40.00% | 60.00% | 80.00% |

## Discussion

### ARB from stranded cetaceans

Most of the bacteria isolated from stranded dolphins were previously reported to cause zoonotic infections, such as *Enterobacter* spp. and *Klebsiella* spp., which belong to the ESKAPE-pathogens (i.e., *Enterococcus faecium, Staphylococcus aureus, Klebsiella pneumoniae, Acinetobacter baumannii, Pseudomonas aeruginosa* and *Enterobacter* species) and are described as the leading cause of resistant nosocomial infections.

*Clostridium perfringens, Actinobacillus delphinicola, A. scotiae, Brucella* spp., *Leptospira* spp., and extended-spectrum *β* -lactamases (ESBL) *Escherichia coli* were also reported to cause infections in marine mammals [40–43]. Buck et al. (1991) isolated *E. coli*, *M. morganii* and *P. mirabilis* from the integumentary system of beluga whale and bottlenose dolphin [20]. Meanwhile, pathogens such as *E. coli,* together with *C. freundii*, *E. aerogenes, K. pneumoniae* and *Klebsiella* spp., were also isolated from the respiratory system of beluga whale. Lastly, *Providencia* spp. were also isolated from the digestive system of beluga whale and bottlenose dolphin. MDR such as ESBL *E. coli,* were isolated from stranded cetaceans [9; 13; 44].

Prior to this study, antibiotic susceptibilities of bacteria isolated from stranded cetaceans were reported, wherein, more than half of the isolates showed single or multiple resistances to the tested antibiotics [9]. The presence of zoonotic pathogens (and associated antibiotic resistances) in these cetaceans indicate biological pollution and presence of antibiotic resistance in their habitats [44–48]

Overall, 5.81% of the bacteria isolated from cetaceans had MAR indices greater than 0.2, which indicates that the environment has a high enteric disease potential [9; 37]. The prevalence of MDR bacteria and genes may be due to the widespread use of antibiotics in humans and livestock, exposure of the cetaceans to bacteria from contaminated sites that may be highly polluted with antibiotics (i.e., domestic, industrial and hospital sewage effluents, water-treatment facilities, livestock production), and exposure of the cetaceans themselves to several antibiotics through medical management of animals (e.g., during stranding events) that may drain to the ocean as runoff, resulting to elevated natural background levels of antibiotic resistance in the marine environment [43; 49–54].

### Prevalence of acquired resistance phenotype from stranded cetaceans

The bacterial isolates have acquired intrinsic resistances to penicillins (e.g., penicillins (e.g., ampicillin), penicillins + β-lactamase inhibitors (e.g., amoxicillin-clavulanic acid) and anti-pseudomonal penicillins + β-lactamase inhibitors antibiotic (e.g., piperacillin-tazobactam), *β* -lactamase (non-extended spectrum cephalosporin (1^st^ and 2^nd^ generation cephalosporins) (e.g., cefuroxime), cephamycin (e.g., cefoxitin), extended-spectrum cephalosporins (3^rd^ and 4^th^ generation cephalosporins) (e.g., ceftazidime, ceftriaxone and cefepime), fluoroquinolones (e.g., ciprofloxacin), polymyxins (e.g., colistin) and folate pathway inhibitors (e.g., trimethoprim-sulphamethoxazole). ‘Multiresistance’ refers to (veterinary) bacteria with acquired resistance properties, and their intrinsic resistance to antimicrobial agents is excluded from MDR definitions [55–56]. MDR veterinary pathogens are further categorized into XDR and PDR bacteria in order to determine the true clinical impact of AMR in therapeutics [38; 55–56]. In this study, 25.80% isolates were confirmed to be either MDR or XDR, which is a contrast to some recent studies that have reported a high proportion of MDR bacteria in the animals tested. For example, Vale et al. (2021) reported that 66.6% (26/39) of *E. coli* isolates from rescued seals (i.e., harbor seals (*Phoca vitulina*) and two grey seals (*H. grypus*) were MDR [12]. Also, Grünzweil et al. (2021) reported that 71% of the ESBL-and *AmpC*-producing Enterobacterales isolates from 148 stranded marine mammals (i.e., California sea lions (*Zalophus californianus*), northern elephant seals (*Mirounga angustirostris*), harbor seals (*P. vitulina*), northern fur seals (*Callorhinus ursinus*), Guadalupe fur seal (*Arctocephalus townsendi*), long-beaked common dolphin (*Delphinus capensis*), and pygmy sperm whale (*K. breviceps*), exhibited MDR phenotype [13]. Despite the low prevalence of MDR and XDR bacteria observed in this current study, the findings indicate biological pollution in the marine environment as well as widespread presence of diverse ARGs that may potentially increase the community acquired resistances [9; 53].

### ARGs from stranded cetaceans

The presence of AMR genes is parallel to the resistance phenotypes exhibited by the *Enterobacteriaceae* isolates [31–35; 57–59]. Among the resistance genes targeted in this study, β-lactamases (*bla*_AmpC_ and *bla*_SHV_), plasmid-mediated-colistin resistance gene (*mcr-1*) and sulfonamide resistance gene (*sul*2) have been detected in seven out of eight isolates that exhibited acquired resistance phenotype. These resistance genes are known to be generally acquired via mobile genetic elements [60–68]. For instance, β-lactamases have diversified in response to the clinical use of β-lactams and are the primary cause of resistance to β-lactams (penicillin, cephalosporins, carbapenems etc.) among members of the family *Enterobacteriaceae* [64].

Aside from β-lactamases, sulfonamide resistance has been observed in Gram-negative bacteria isolated from humans and animals, and it is mediated via acquisition of alternative dihydropteroate synthase gene (*sul*) with several variants (e.g., *sul*1*, sul*2 and *sul*3), causing the bacteria to develop low affinity for sulfonamides [61–62; 69]. It is believed that *sul*2, which is detected in this study, is usually located in small nonconjugative plasmids or large transmissible multiresistance plasmids [60–62]. The emergence and spread of plasmid-mediated colistin-resistance gene (*mcr-1*) in livestock, food, and humans, pose a threat on the effectiveness of colistin, the last-resort antibiotic used to treat MDR bacteria-related infections caused by carbapenem-resistant *Enterobacteriaceae* [65–68].

The low prevalence of ARGs in the isolates is in congruence with a previous study that also reported lower abundances of ARGs in marine aquatic animals compared to freshwater aquatic animals [70]. Freshwater environments are said to easily receive antibiotic residues as compared to marine waters owing to their proximity to sites highly contaminated with antibiotics, thus freshwater animals are expected to harbor more ARGs [70]. Recent studies reported the prevalence of ARGs in marine mammals (e.g., rescued seals and stranded marine mammals) and found that the bacteria isolated from these marine animals possess β-lactamase encoding genes, *bla*_OXA−1_, *bla*_TEM−1,_ *bla*_CMY,_ *bla*_SHV-33,_ *bla*_CTX-M-15_, *bla*_OXA-15_, *bla*_SHV-11_ and *bla*_DHA-11._ Meanwhile, the most prevalent non-*β*-lactamase gene was *sul*2, followed by *str*A, *str*B and *tet*(A) [12–13].

### Cetacean characteristics and antibiotic resistance profiles

The cetacean species sampled in this study generally inhabit deep or offshore areas, where upwelling occurs. These include short-finned pilot whale (*Globicephala macrorhynchus*), striped dolphins (*S. coeruleoalba*), pygmy sperm whale (*K. breviceps*), Fraser’s dolphin (*L. hosei*), melon-headed whale (*P. electra*) and Bryde’s whale (*B. edeni*) [71–76]. Some species such as bottlenose dolphins (i.e., Indo-pacific bottlenose dolphins (*T. aduncus*) prefer coastal waters [76–77]. Few species, such as Risso’s dolphin (*G. griseus*), are known widespread species which occur in all habitats from coastal to oceanic waters [78]. In this study, all isolates with higher MAR index values of > 0.2 were isolated from cetacean species that inhabit offshore areas, such as pygmy sperm whale (*K. breviceps*) and short-finned pilot whale (*G. macrorhynchus*), with 27.27% and 50% of the bacteria isolated from each species respectively. This may be due their physiology which requires a regular need to return to the air-water interface for several minutes to breathe, in order to set biochemical stage of successful subsequent dives, thus exposing themselves to several forms of pollution that eventually reach them from the nearby coast [79–80].

### Animal tissues as niche for the dissemination of antibiotic resistance in water environments

Some studies reported that microbiota in animal tissues or body parts, such as those in skin microbiota, may contribute to the proliferation and propagation of antibiotic resistance and ARGs in water environment [8; 70]. In this study, source/s of the swab samples were taken into consideration and it was found that more bacteria with MAR index values of > 0.2 were isolated from tissues in contact with the environment such as in the anus and blowhole, with 40.00% and 15.38% percentage of the isolates, respectively. This is in agreement with the previous report of Rose et al., (2009) wherein tissues (e.g., oral, cloacal and blowhole) of animals that have direct contact to water had higher incidence of bacteria with resistance to multiple antibiotics as compared to internal tissues (e.g., spleen and thorax) and fecal samples [8]. This suggests that tissues in direct contact with the environment (e.g., skin) may be an important niche for the dissemination of ARGS in the environment [70].

## Conclusion

This study investigated *Enterobacteriaceae* isolated from nineteen stranded cetaceans for phenotypic and genotypic resistance to select antibiotics. The isolated bacteria belong to the genera *Escherichia, Enterobacter, Klebsiella, Citrobacter, Morganella, Pantoea,* and *Providencia*. Overall, 35/86 (40.70%) of the isolates exhibited phenotypic resistance to cephalosporins, polymyxins, folate-pathway inhibitors, and ampicillin, while resistance against amoxicillin-clavulanic acid, piperacillin, and imipenem was relatively low. Furthermore, isolates were characterized as multidrug-resistant (MDR) (2.33%) and extensively drug-resistant (XDR) (38.37%). As for genotypic resistance, six out of seven target antibiotic resistance genes (ARGs) responsible for resistance to ampicillins, cephalosporins, polymyxins, and sulfonamides were detected in nearly half of the isolates with acquired resistance. The cetacean species included in the study inhabit deep waters but frequently surface to breathe, potentially exposing themselves to various forms of pollution. This exposure may contribute to the proliferation and dissemination of antibiotic resistance, particularly in tissues and body parts that come into direct contact with the environment, such as the oral cavity, cloaca, and blowhole.

Overall, the findings of study suggest the wide distribution of antibiotics and ARGs in the marine environment and supports the potential contribution of cetaceans to the spread of antibiotic resistance in their habitats. Further research and mitigation efforts are necessary to better understand and mitigate the impact of antibiotic resistance in marine environments.

## Acknowledgments

We thank the Philippine Marine Mammal Stranding Network (PMMSN) and the Bureau of Fisheries and Aquatic Resources (BFAR) for the nationwide cetacean stranding response. We are also grateful to Dr. Wei-Cheng Yang for the valuable inputs and review of the manuscript. This project is funded by Ocean Park Conservation Foundation Hong Kong Project No. MM02_1819.

## Authors’ contributions

RMDV, MCMO, WLR, MATS, and LVA conceptualized and designed the study. RMDV, JAAC and MCMO performed the laboratory procedures. WLR provided the facility for, and supervised the VITEK testing. LVA coordinated the stranding response with RMDV, MCMO, and JAAC. MCMO, WLR, MATS, and LVA received the funding for the study. RMDV, MCMO, WLR, and MATS analyzed data. All authors contributed to the writing and approved the final version of the manuscript.

## Supporting Information

**S1 Table 1. Bacteria isolated from stranded cetaceans in the Philippines from 2017-2019**

**S2 Table 2. Phenotypic antibiotic resistance patterns of *Enterobacteriaceae* isolated from stranded cetaceans**

**S3 Table 3. Antibiotic resistance profiles of *Enterobacteriaceae* isolates from stranded cetaceans**

**S4 Table 4. Incidence of multiple antibiotic resistances among *Enterobacteriaceae* isolated from different cetacean species.**

**S5 Table 5. Incidence of multiple antibiotic resistances among *Enterobacteriaceae* isolated from different source samples.**

**S6 Table 6. Incidence of multiple antibiotic resistances among *Enterobacteriaceae* isolated from different administrative regions.**

**S1 Fig 1 *bla*_CTX-M_ detection (a. Cefuroxime-resistant *Enterobacteriaceae* isolates & b. Cephalosporin-resistant and penicillin-resistant *Enterobacteriaceae* isolates)**

**S2 Fig 2. *bla*_TEM_ detection (a. Cefuroxime-resistant *Enterobacteriaceae* isolates & b. Cephalosporin-resistant and penicillin-resistant *Enterobacteriaceae* isolates)**

**S3 Fig 3. *bla*_SHV_ detection (a. Cefuroxime-resistant *Enterobacteriaceae* isolates & b. Cephalosporin-resistant and penicillin-resistant *Enterobacteriaceae* isolates)**

**S4 Fig. 4. *bla*_AMP-C_ detection (a-c. Penicillin-resistant *Enterobacteriaceae* isolate)**

**S5 Fig. 5. *bla*_KPC_ detection**

**S6 Fig. 6. mcr-1 detection**

**S7 Fig. 7. *sul*I detection**

**S8 Fig. 8. *sul*II detection**

